# Germplasm of Brazilian winter squash (*Cucurbita moschata* D.) displays vast genetic variability, allowing identification of promising genotypes for agro-morphological traits

**DOI:** 10.1101/2020.03.04.977454

**Authors:** Ronaldo Silva Gomes, Ronaldo Machado Júnior, Cleverson Freitas de Almeida, Rafael Ravaneli Chagas, Rebeca Lourenço de Oliveira, Fabio Teixeira Delazari, Derly José Henriques da Silva

## Abstract

Winter squash fruits (*Cucurbita moschata* D.) are among the best sources of vitamin A precursors and constitute sources of bioactive components such as phenolic compounds and flavonoids. Approximately 70% of *C. moschata* seed oil is made up of unsaturated fatty acids, with high levels of monounsaturated fatty acids and components such as vitamin E and carotenoids, which represent a promising nutritional aspect in the production of this vegetable. *C. moschata* germplasm expresses high genetic variability, especially in Brazil. We assessed 91 *C. moschata* accessions, from different regions of Brazil, and maintained at the UFV Vegetable Germplasm Bank, to identify early-flowering accessions with high levels of carotenoids in the fruit pulp and high yields of seed and seed oil. Results showed that the accessions have high variability in the number and mass of seeds per fruit, number of accumulated degree-days for flowering, total carotenoid content, and fruit productivity, which allowed selection for considerable gains in these characteristics. Analysis of the correlation between these characteristics provided information that will assist in selection to improve this crop. Cluster analysis resulted in the formation of 16 groups, confirming the variability of the accessions. *Per se* analysis identified accessions BGH-6749, BGH-5639, and BGH-219 as those with the earliest flowering. Accessions BGH-5455A and BGH-5598A had the highest carotenoid content, with averages greater than 170.00 μg g^-1^ of fresh mass. With a productivity of 0.13 t ha^-1^, accessions BGH-5485A, BGH-4610A, and BGH-5472A were the most promising for seed oil production. These last two accessions corresponded to those with higher seed productivity, averaging 0.58 and 0.54 t ha^-1^, respectively. This study confirms the high potential of this germplasm for use in breeding for promotion of earlier flowering and increase in total carotenoid content of the fruit pulp and in seed and seed oil productivity.

## Introduction

Winter squash (*Cucurbita moschata* D.) is one of the vegetables of greater socio-economic importance in the *Cucurbita* genus, largely due to the high nutritional value of its fruits and seeds. The pulp of its fruits comprises an important source of carotenoids such as *β*-carotene, the precursor of greater pro-vitamin A activity [1, 2, 3], in addition to vitamins such as B2, C, and E [4, 5]. The pulp is also an excellent source of minerals such as K, Ca, P, Mg, and Cu [6, 7]. The socio-economic importance of *C. moschata* is also linked to the high volume and value of its production. It is estimated that, together with other cucurbits such as *C. pepo* and *C. maxima*, the cultivated area and the world production of this vegetable in 2017 were approximately 2 million hectares and 25 million tons, respectively [8], most of it concentrated in China and India. In Brazil this crop is of high socio-economic importance, with a cultivated area of approximately 90 thousand hectares, an estimated production of more than 40 thousand tons / year and an annual production value of around R$ 1.5 million [9].

*C. moschata* brings together characteristics that are fundamental in biofortification programmes, such as high productivity and profitability potentials, high efficiency in reducing micronutrient deficiencies in humans, and good acceptability with producers and consumers in its growing regions [10]. This has caused the vegetable to be chosen as a strategic crop for breeding programmes promoting biofortification, such as the Brazilian biofortification programme (BioFORT), led by Embrapa, which aims for biofortification in vitamin A precursors [11]. The potential of *C. moschata* for the production of edible seed oil is also a promising aspect of this crop. Constituted of about 70% unsaturated fatty acids with a high content of monounsaturated fatty acid [12, 13], *C. moschata* seed oil is a good substitute for other lipid sources with higher saturated fatty acid contents. The oil is also rich in bioactive components such as vitamin E and carotenoids [4], which are important antioxidants in the human diet, in addition to providing protection to the oil against oxidative processes. In addition, the cultivation of this species is commonly based on a production system of low-technological level [14], making this crop fundamental for ensuring healthier diets and promoting food security in the regions where it is grown, particularly in less developed regions and in the context of family-based farming.

Associated with its socio-economic importance, *C. moschata* germplasm commonly expresses high genetic variability in all regions where it occurs [15, 16, 17], especially in Brazil [18, 19, 20]. Archaeological evidence indicates that this species was present in Latin America prior to colonisation, and appears to have already been an important component in the diet of the native peoples living there [21, 22, 23]. Currently, the variability of this vegetable in Brazil is closely tied to the human populations involved in its cultivation, who are predominantly family-based farmers. The selection practised over time by these populations, associated with the exchange of seeds between them, and the natural occurrence of hybridization in the germplasm of this species has contributed to the increase in its variability. The high variability displayed by *C. moschata* for agronomic, nutritional and bioactive characteristics and the intercrossability of the *Cucurbita* species has enabled the transfer of these characteristics from *C. moschata* to other species of this genus [24, 25, 26, 27]. This is something strategic and may aid the worldwide cultivation of species of the *Cucurbita* genus.

The usefulness of plant germplasm conserved in banks depends on the amount and quality of information associated with it, which highlights the importance of its proper evaluation. On the other hand, the high volume of germplasm and limitations in resources and area available for the establishment of field trials commonly make its assessment difficult. In view of this, the FAO’s Second Global Action Plan for Plant Genetic Resources for Food and Agriculture sets out guidelines that provide greater efficiency in the conservation and use of plant germplasm [28]. This is essential information for the management and use of germplasm [14, 29, 30, 31]. Evaluation of the germplasm maintained in banks makes it possible to estimate the magnitude of the genetic and statistical parameters of characteristics of interest, which can provide information on the nature of variability observed for these traits, in addition to elucidating which characteristics or groups of characteristics most contribute to germplasm variability. From this assessment, it is also possible to assess the association between the characteristics evaluated. Together, the information obtained from these assessments is essential for the optimisation of the use and management of plant germplasm.

The UFV Vegetable Germplasm Bank (BGH-UFV) maintains more than 350 accessions of *C. moschata*, comprising one of the largest collections of this species in the country [32]. This bank continually carries out work on the characterisation and evaluation of this germplasm [33], which has allowed the identification of sources of resistance for important phyto-pathogenic agents of this crop [34], and for its improvement in terms of production [20] and nutritional aspects of fruits and seed oil [12, 35]. The potential of this germplasm as a source of genes for the improvement of this crop, along with the possibility of elucidating the genetic mechanisms linked to important production parameters, justifies the continuation of studies on its assessment and use.

This study therefore aimed to: a) agro-morphologically assess some of the *C. moschata* accessions maintained by BGH-UFV, b) analyse the genetic relationships of the agro-morphological characteristics, and c) analyse the agro-morphological variability, with a view to identifying earlier-flowering genotypes, genotypes with high total levels of carotenoids in the fruit pulp, and those with high potential for seed and seed oil productivity.

## MATERIALS AND METHODS

### Origin of germplasm and preparation of seedlings

In this study, we assessed 95 genotypes, which comprised 91 accessions of *C. moschata* maintained in the BGH-UFV, and four control genotypes (Table 1). The accessions came from different regions of Brazil [33], and consisted, for the most part, of samples collected from family-based farmers, who commonly perform the selection of genotypes and the conservation of their seeds.

**Table 1.** Origin of the of the *C. moschata* accessions and controls assessed in this study and maintained by BGH-UFV

| Origem | Number of accessions |
| --- | --- |
| Bahia | 6 |
| São Paulo | 39 |
| Minas Gerais | 15 |
| Distrito Federal | 17 |
| Espírito Santo | 3 |
| Goiás | 4 |
| Paraná | 2 |
| Rio de Janeiro | 2 |
| Rio Grande do Norte | 2 |
| Santa Catarina | 1 |
| Controls |  |
| Jabras- Isla Sementes |  |
| Jacarezinho- Embrapa Hortaliças |  |
| Maranhão- Embrapa Hortaliças |  |
| Tetsukabuto- Sakata Sementes |  |

Seedlings were produced in a 72-cell expanded-polystyrene tray containing commercial substrate. Seedling transplantation and cultural treatments were carried out according to local recommendations for the cultivation of pumpkins [36].

### Experiment location and experimental design

The experiment was carried out from January to July 2016, at “Horta Velha” (200 45’14 ‘’ S, 420 52’53 ‘’ W and 648.74 m), an experimental unit of the Agronomy Department of the Federal University of Viçosa, Viçosa-MG, Brazil.

The experiment was arranged in Federer’s augmented block design [37], with five repetitions for each control. The four controls, also called common treatments, were randomly distributed in each of the five blocks, and the 91 accessions, called regular treatments, were randomly assigned to all blocks. A spacing of 3×3 m between plants and rows was adopted, which resulted in a stand of 1,111 plants ha^-1^. Each plot consisted of five plants, of which the three central plants were considered for evaluation.

### Assessments of agro-morphological aspects, total carotenoid content of fruit pulp, and seed and seed oil yields

For the assessment involving multi-categorical characteristics, we adopted the morphological descriptors suggested by Bioversity and the European Cooperative Program for Plant Genetic Resources (ECPPGR), plus some additional descriptors.

These descriptors comprised agro-morphological characteristics of plants, fruits and seeds, as well as the phytosanitary aspect of plants (supplementary table 1). Assessment was also performed based on agronomic characteristics, the total carotenoid content of fruit pulp and the yields of seed and seed oil (Table 2).

**Table 2.** Descriptors involving agronomic aspects, the total *carotenoid* content of fruit pulp and yields of seed and seed oil used in the assessment of the *C. moschata* germplasm maintained by BGH-UFV

| Phase/organ | Descriptors |
| --- | --- |
| Reproductive phase | Degree-days accumulated for flowering (DDF). |
| Fruit | Number of fruits per plant (NFP), average mass of fruits (MF), productivity of fruits (PF), height of fruit (HF), diameter of fruit (DF), thickness of fruit peel (TFP), resistance of fruit peel (RFP), resistance of fruit pulp (RP), thickness of fruit pulp (PT), diameter of internal cavity of fruit (DIC), total content of fruit pulp carotenoids (TC) and the lutein content of fruit pulp (L). |
| Seed | Number of seeds per plant (NSF), mass of seeds per fruit (MSF), relationship between the masses of seeds and fruit (MS/F), mass of one hundred seeds (MOH), productivity of seeds (PS), seed thickness (ST), seed length (SL), and seed width (SW). |
| Seed oil | Seed oil content (SOC) and seed oil productivity (SOP). |

The estimates of the total carotenoid and lutein contents of the fruit pulp were obtained based on colorimetric parameters. For this, the fruit pulp colour was characterised with the aid of a manual tri-stimulus colorimeter, Color Reader CR-10 Konica Minolta, based on parameters related to luminosity (L), and contribution of red (a) and yellow (b). The characterisation was carried out from different regions in the fruits (region facing the sun, region facing the soil, region of the peduncle and floral insertion), from one fruit of each plant in the useful area of the plot. These estimates were obtained using the equations proposed by [38], described below:

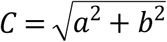

TC= 6.1226 + 1.7106 a

L= −6.3743 + 0.2818 C

Where:

TC corresponds to the total *carotenoid* content of *the* fruit pulp (µg g^-1^ of fresh pulp mass); a corresponds to the contribution of red to the colour of the fruit pulp; L corresponds to the lutein content of the fruit pulp (µg g^-1^ of fresh pulp mass); and C corresponds to the saturation or chroma of the fruit pulp.

The extraction of oil from seeds was carried out by means of cold pressing, with the aid of a 30-ton-capacity press, with the necessary adaptations for pressing. For this, the seeds were previously dried in a forced-air-circulation oven for 72 hours, at 23°C. For standardization of the process, 50 g seed samples were weighed from each access and all samples were equally pressed for approximately 10 minutes.

### Estimation of genotypic values, components of variance and genetic-statistical parameters

Phenotypic data were analysed using maximum restricted likelihood (REML) procedures and the best unbiased prediction (BLUP). These procedures were carried out with the aid of the R program, using the “lme4” package [39]. The estimates of variance components were obtained from this first procedure, while the genotypic values of accessions (BLUPS) and controls (BLUES), were obtained from the BLUP procedure. All estimates were obtained based on the following model:

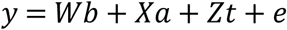

In which:

y corresponds to the phenotypic data vector;

b corresponds to the vector comprising the effect of block, assumed to be random;

a corresponds to the vector comprising the effect of accessions, assumed to be random:

t corresponds to the vector comprising the effect of controls, assumed to be fixed: and e corresponds to the error vector.

The letters W, X and Z correspond to the incidence matrices of parameters b, a, and t, respectively, with the data vector y.

The estimates of the variance components comprised the phenotypic (σ^2^_*p*_), genotypic (σ^2^_*g*_), and residual (σ^2^) variances, and the variance associated with the block effect (σ^2^_*b*_). The genetic-statistical parameters comprised the broad sense heritability (*h*^2^), the selection accuracy (*A*), selection gain (*SG*), the phenotypic mean of the characteristics (*μ*), and the genotypic (CV_*g*_ %), phenotypic (CV_*P*_ %), and residual (CV*r* %) coefficients of variance. These were obtained from the following estimators: *h*^2^ = 1-(*Pev*/2σ^2^_*g*_), where *Pev* corresponds to the prediction of errors variance [40]; 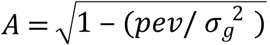 ; *GS* = *h*^*2*^. *DS*, where *DS* corresponds to the selection differential, estimated based on the average of the top 15% most promising accessions; CV_*g*_ % = (σ^2^_*g*_/*μ*) x 100; CV_*p*_ % = (σ^2^_*p*_/*μ*) x 100; e CV*r* % = (σ^2^_*g*_/*μ*) x 100.

### Correlation analysis

This analysis was performed based on the matrix of genetic correlations, obtained based on the following estimator:

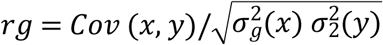

In which;

*Cov* (x, y), corresponds to the genetic covariance between two variables X and Y, and σ^2^_*g*_(x) and σ^2^_*g*_ (y) correspond to the genetic variances of variables X and Y, respectively.

The correlations were analysed using a procedure known as *correlation network*, which allows, based on a specific function, the analysis of all relationships between the variables under study. This procedure also allows the direction and magnitude of the correlations to be distinguished. The direction is denoted by colours; a dark green colour is used for the lines that connect positively-correlated variables, and a red colour for the lines that connect negatively-correlated variables. The magnitude of the correlations is denoted by the thickness of the lines connecting the variables; the thicker the line, the greater the correlation. The significance of the correlations was analysed using Mantel’s Z test at 1 and 5% probability. The correlation analysis was performed with the aid of the Genes program [41].

### Analysis of variability and clustering

The analysis of variability was carried out using both quantitative and qualitative information. For quantitative data, the distance matrix between the genotypes was obtained from the BLUPS estimates in the case of accessions, and BLUES in the case of the controls, and were estimated based on the negative average Euclidean distance, with data standardization.

The matrix was obtained from *negDistMat*, a function of the APCluster package [42] implemented in the R program, version 3.5.1 (R Development Core Team, Vienna, AT). The distances d (x; y) between the accession pairs, exemplified here as any two accessions x (x_1_, …, x_n_) and y (y_1_, …, y_n_), were estimated based on the following equation:

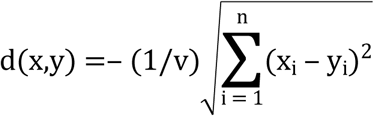

In which v corresponds to the number of quantitative descriptors evaluated.

The distance matrix for the qualitative data was obtained using the arithmetic complement of the simple coincidence index. The variability analysis was performed from a single distance matrix, obtained from the sum of the distance matrices of the quantitative and qualitative data. For the sum of matrices, they were standardised and each received an equal weight in the sum procedure. The variability analysis was performed using the procedure known as the *Affinity propagation* method [43]. The grouping was carried out from 100 independent rounds, aiming to assess the consistency of grouping.

The operation of *Affinity* initially involves the identification, in a set of components, of samples that will function as centres of this set. This method simultaneously considers all the set components as potential centres, i.e. as nodes in an interconnected network. Following the identification of potential centres, messages are transmitted between the set components along the network until a good set of centres and their corresponding groups emerge. The messages exchanged between the components in *Affinity* can be “responsiveness” *r* (*i, k*) and “availability *a* (*i, k*). This first case reflects the accumulated evidence of how appropriate point *k* is to serve as an example for point *i*, considering all other potential examples for this point. The “availability”, in turn, reflects the accumulated evidence of how appropriate it would be for point *i* to choose point *k* as an exemplar, considering the other points for which point *k* can be an exemplar [43]. In the analysis of the present study, the availability was initially established as zero.

### Identification of promising accession groups and *per se* identification of accessions

In order to facilitate the identification of promising groups of accessions for each characteristic, we carried out a grouping of means of the genotypic values corresponding to the groups obtained from the analysis of variability. This was carried out based on Tocher’s grouping of means method. The identification *per se* of the most promising accessions for each trait was carried out by ranking the respective genotypic effects, genetic gain and the new predicted average of the accessions, and the top 15% were considered the most promising accessions.

## RESULTS

### Variance components and genetic-statistical parameters

Estimates of the variance components and the genetic-statistical parameters are presented in table 3. The highest estimates of genotypic variance corresponded to the number of seeds (NSF) and mass of seeds per fruit (MSF), to the degree-days accumulated for flowering (DDF), and to the total carotenoid content of the fruit pulp (TC). Among these variance estimates, only the genotypic variance of DDF was not significant.

**Table 3.** Estimates of variance components and genetic-statistical parameters of agronomic aspects, the total carotenoid content of fruit pulp, and of the yields of seeds and seed oil of the *C. moschata* germplasm accessed in this study and maintained by the BGH-UFV

| Vegetative trait |  |  |  |  |  |  |  |  |  |  |  |  |
| --- | --- | --- | --- | --- | --- | --- | --- | --- | --- | --- | --- | --- |
| Traits | $\sigma_p$ | $\sigma_g$ | $\sigma$ | $\sigma_b$ | $A$ | $h^2$ | $SG$ | Range | $\mu$ | $CV_g$ % | $CV_p$ % | $CV_r$ % |
| DDF | 10781.493 | 6385.892<sup>ns</sup> | 3909.203 | 486.397800 | 0.725 | 0.525 | -92.947 | 120.0- 820.4 | 606.642 | 13.172 | 17.116 | 10.306 |
| Fruit traits |  |  |  |  |  |  |  |  |  |  |  |  |
| Traits | $\sigma_p$ | $\sigma_g$ | $\sigma$ | $\sigma_b$ | $A$ | $h^2$ | $SG$ | Range | $\mu$ | $CV_g$ % | $CV_p$ % | $CV_r$ % |
| NFP | 8.724 | 3.583<sup>ns</sup> | 4.614 | 0.527 | 0.655 | 0.429 | 2.303 | 1- 15 | 4.783 | 39.575 | 61.752 | 44.909 |
| MF | 2.738 | 2.373** | 0.364 | 0.000 | 0.841 | 0.707 | 2.189 | 0.45-10.0 | 2.735 | 56.323 | 60.500 | 22.059 |
| PF | 73.954 | 38.598* | 29.279 | 6.076 | 0.704 | 0.495 | 7.817 | 0.7- 44.6 | 12.946 | 47.989 | 66.427 | 41.796 |
| TC | 387.206 | 362.902** | 24.303 | 0.000 | 0.880 | 0.774 | 20.426 | 43.4- 187.2 | 65.763 | 28.967 | 29.921 | 7.496 |
| Seed and oil traits |  |  |  |  |  |  |  |  |  |  |  |  |
| Traits | $\sigma_p$ | $\sigma_g$ | $\sigma$ | $\sigma_b$ | $A$ | $h^2$ | $SG$ | Range | $\mu$ | $CV_g$ % | $CV_p$ % | $CV_r$ % |
| NSF | 25274.617 | 20784.317** | 2817.703 | 1672.597 | 0.844 | 0.712 | 167.873 | 78.6- 805.7 | 454.188 | 31.741 | 35.003 | 11.685 |
| MSF | 490.881 | 465.357** | 16.141 | 9.382 | 0.899 | 0.808 | 27.428 | 4.4- 119.3 | 51.929 | 41.541 | 42.665 | 7.736 |
| MS/F | 0.000142523 | 0.000125343** | 0.000015091 | 0.000002089 | 0.854 | 0.729 | 0.015 | 0.00- 0.05 | 0.023 | 48.676 | 51.905 | 16.890 |
| MOHS | 7.395 | 4.391* | 3.003 | 0.000 | 0.721 | 0.519 | 2.210 | 6.3- 23.6 | 11.701 | 17.908 | 23.240 | 14.809 |
| PS | 0.042 | 0.019<sup>ns</sup> | 0.016 | 0.006 | 0.694 | 0.481 | 0.187 | 0.01- 0.9 | 0.269 | 51.241 | 76.185 | 47.022 |
| SOC | 13.010 | 0.462<sup>ns</sup> | 11.822 | 0.725 | 0.512 | 0.262 | 1.254 | 28.50- 54.4 | 18.516 | 3.670 | 19.480 | 18.569 |
| SOP | 0.001743 | 0.000172<sup>ns</sup> | 0.001300 | 0.000 | 0.540 | 0.291 | 0.072 | 0.004- 0.40 | 0.050 | 26.037 | 83.498 | 72.111 |

Degree-days accumulated for flowering (DDF), number of fruits per plant (NFP), average mass of fruits (MF), productivity of fruits (PF), total carotenoid content of fruit pulp (TC), number of seeds per fruit (NSP), mass of seeds per fruit (MSF), relationship between the masses of seeds and fruit (MS/F), mass of one hundred seeds (MOHS), productivity of seeds (PS), seed oil content (SOC), and seed oil productivity (SOP). Components of variance involving phenotypic ( $\sigma_p$ ), genotypic ( $\sigma_g$ ), residual ( $\sigma$ ) and the variance associated to block effect ( $\sigma_b$ ). Genetic-statistical parameters involving accuracy ( $A$ ), broad sense heritability ( $h^2$ ), selection gain ( $SG$ ), average ( $\mu$ ), coefficients of genotypic ( $CV_g$ %), phenotypic ( $CV_p$ %) and residual variation ( $CV_r$ %). *ns* not significant; \*\*, \* significant at $p < 0.01$ and $0.05$ , respectively by the likelihood ratio test.

The greatest contributions from the genotypic variance (σ^2^_*g*_), to the phenotypic (σ^2^_*p*_) also corresponded to the characteristics of seeds, namely MSF (94.80%) and NSF (82.23%). A significant contribution was also observed for TC (93.72%) and DDF (59.23%), as shown in table 3. The residual variance had a reduced contribution to the phenotypic variance for most of the characteristics. As can also be seen in table 3, most of the characteristics displayed high estimates for the selection accuracy (*A*). The heritability estimate of DDF was 0.525; and 0.495 and 0.774 for PF and TC, respectively. PS expressed broad sense heritability of 0.481 and SOP 0.291. The selection gain estimate for DDF was -92.947, and was 7,817 t ha^-1^ and 20.426 µg g^-1^ of fresh pulp mass for PF and TC, respectively. The selection gains for PS and SOP were 0.187 and 0.072 t ha^-1^, respectively (Table 3).

Phenotypic amplitude for DDF between accessions was 120.0 to 820.4 and the average was 606. 642 (Table 3). The amplitude for PF was 44.6 to 0.7 t ha^-1^ and the average 12.946 t ha^-1^. The amplitude for TC was 43.4 to 187.2 µg g^-1^ of fresh pulp mass and the average was 65.763 µg g^-1^ of fresh pulp mass. The amplitude for PS was 0.01 to 0.9 t ha^-1^ and the average was 0.269 t ha^-1^. The phenotypic and average amplitudes of the accessions for SOP were 0.004 to 0.40 t ha^-1^ and 0.050 t ha^-1^, respectively (Table 3). The greatest amplitude for the coefficient of genotypic variation (CV_*g*_%) was observed between the characteristics SOC and MF, while for the coefficient of phenotypic variation (CV_*P*_%), the greatest amplitude was observed between DDF and SOP. The estimates of residual variation coefficient ranged from 7.502 to 71.582 for TC and SOP, respectively (Table 3).

### Genotypic correlation

The genotypic correlation network of agronomic aspects, the total carotenoid content of the fruit pulp, and the characteristics of seeds and seed oil are shown in figure 1. Visually, it is possible to observe cohesion of groups involving some of the fruit characteristics and groups involving some of the characteristics of seeds. It is possible to observe cohesion between fruit productivity and other characteristics of this group such as MF, DIC, HF, DF and TPF. As can be inferred from the colour and thickness of lines, this set of variables expressed positive correlations of high magnitudes. The highest correlations in this group correspond to the correlations of fruit yield with MF (0.61) and with DIC (0.54), which were also significant (*p*<0.01) (Figure 1). There was cohesion between the group of variables involved in seed productivity and variables such as MS/F, NSF and MSF. This set of variables expressed positive and high-magnitude correlations, and the correlation of seed productivity with MS/F (0.56), which was significant (*p*<0.01), was the highest in this group. The group involving the mass of one hundred seeds and characteristics of seed dimensions such as SW, ST and SL was also a cohesive group. This group expressed positive correlations, and the correlation of the mass of one hundred seeds with SW (0.62), also significant (*p*<0.01), was the highest in this group (Figure 1).

**Figure 1.**
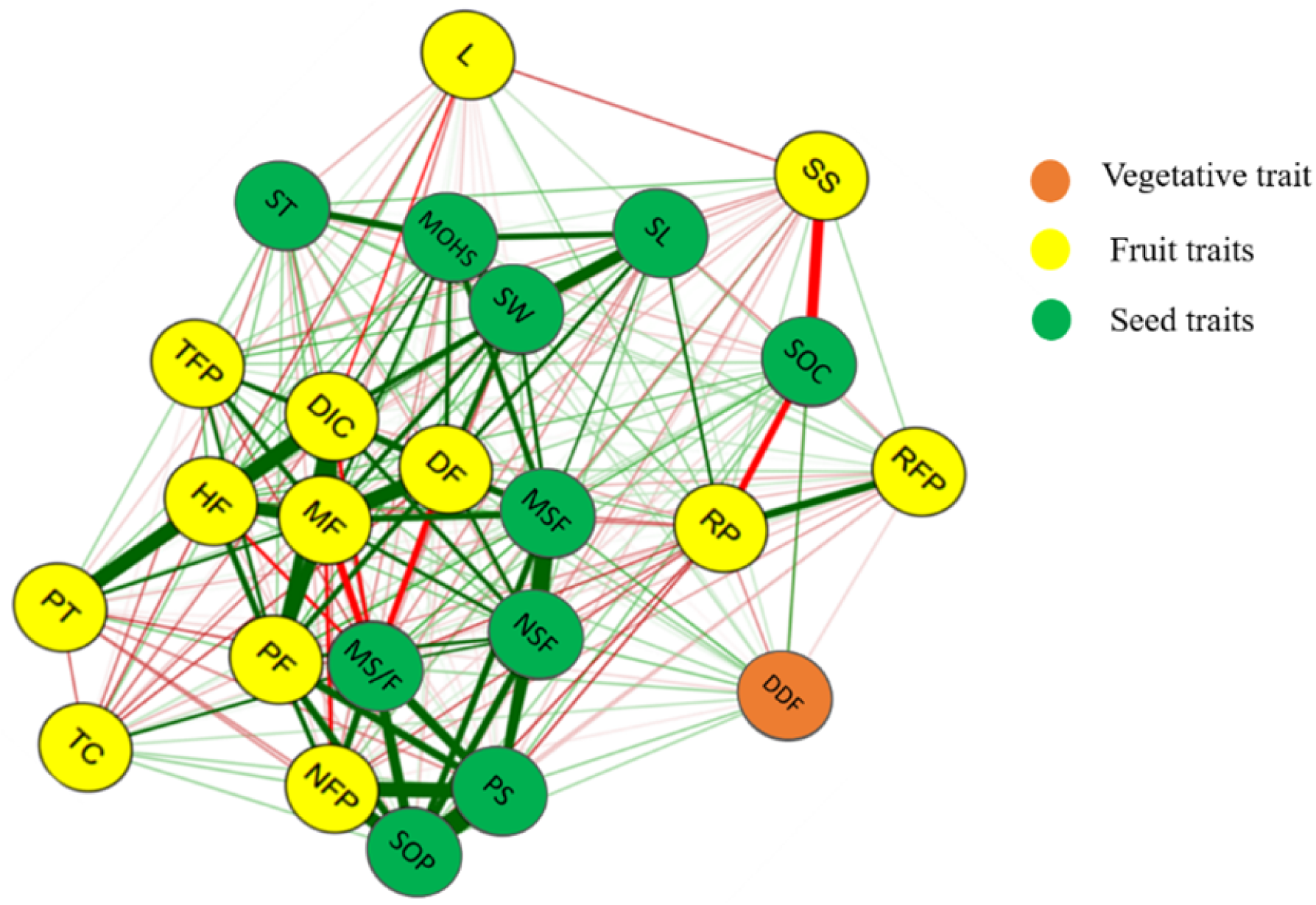
Network of genotypic correlations of agronomic aspects, the total carotenoid content of the fruit pulp, and the characteristics of seeds and seed oil of the *C. moschata* germplasm assessed in this study and maintained by the BGH-UFV. The red and green lines denote positive and negative correlations, respectively. Thicker lines indicate greater magnitudes of correlation while the thinner lines indicate lesser magnitudes. Degree-days accumulated for flowering (DDF), number of fruits per plant (NFP), average mass of fruits (MF), productivity of fruits (PF), height of fruit (HF), diameter of fruit (DF), thickness of fruit peel (TFP), resistance of fruit peel (RFP), resistance of fruit pulp (RP), pulp thickness (PT), diameter of internal cavity of fruit (DIC), soluble solids of fruit pulp (SS), total carotenoids content of fruit pulp (TC), lutein content of fruit pulp (L), mass of seeds per fruit (MSF), productivity of seeds (PS), relationship between the masses of seeds and fruit (MS/F), mass of one hundred seeds (MOHS), seed oil content (SOC), and seed oil productivity (SOP).

### Genetic variability and clustering

Cluster analysis, based on the agro-morphological aspects, the total carotenoid content of the fruit pulp, and the characteristics related to the yields of seed and seed oil of the germplasm, placed the accessions into 16 groups (Table 4).

**Table 4.**
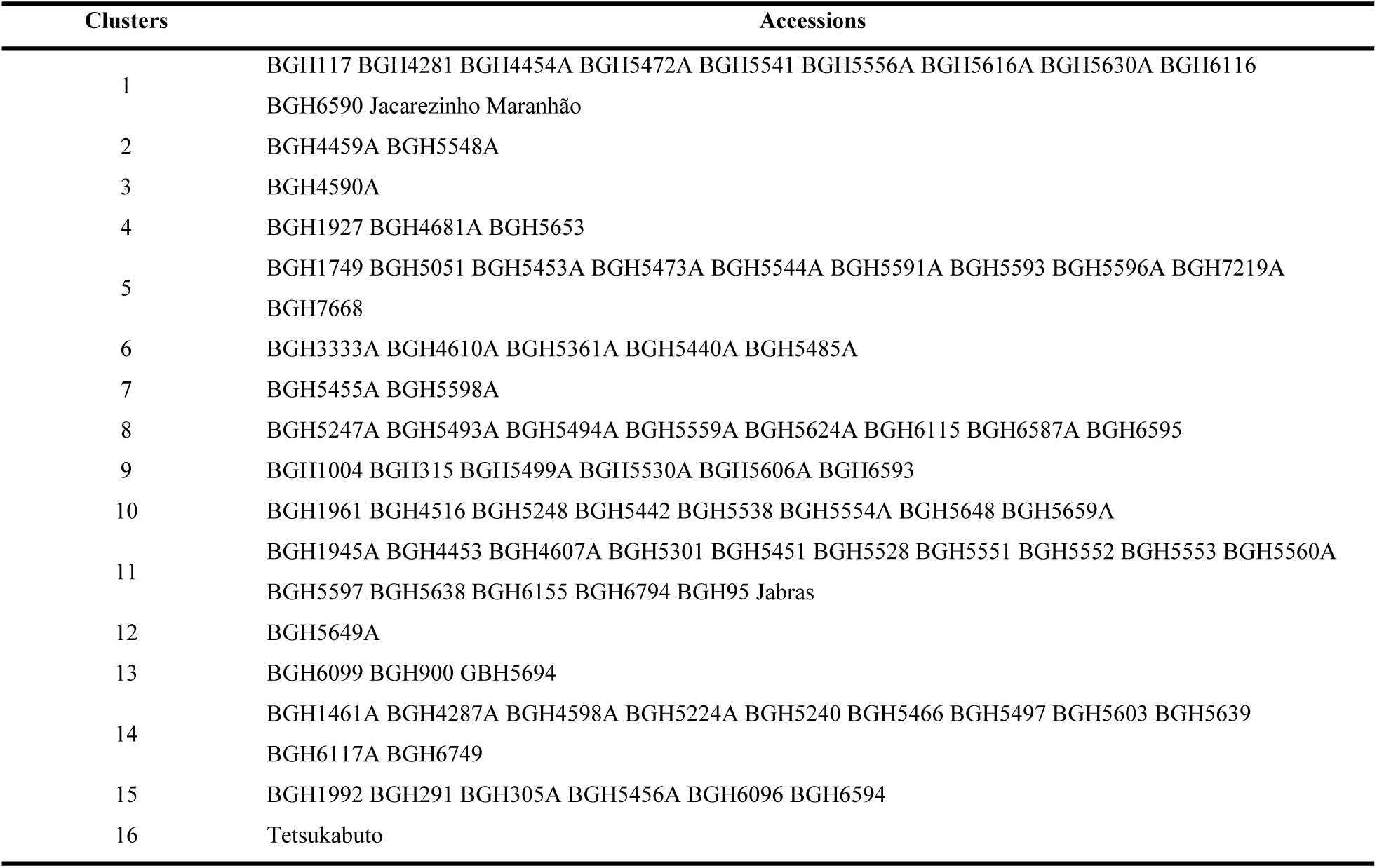
Clustering of the *C. moschata* germplasm assessed in this study and maintained by BGH-UFV, based on agro-morphological aspects, the total carotenoid content of fruit pulp, and the yields of seeds and seed oil

Based on the clustering pattern, high variability was observed between the accessions. Most of the genotypes were grouped in group 11, which comprised 17.58% of the accessions, and the control Jabras, making it the largest group. Group 1, the second largest, contained 13.18% of the accessions and two controls, Jacarezinho and Maranhão. Group 14 was the third largest, in which 12.08% of accessions were grouped. The grouping of genotypes in the other groups did not occur equitably and some of them contained only one genotype (Table 4).

The visual pattern of the clustering in heatmap format showed low similarity between the groups formed, something denoted by the predominance of yellow and orange colouring (Figure 2). Visual analysis of this clustering also shows homogeneity of the distances between groups, denoted by the uniformity of the figure colouring.

**Figure 2.**
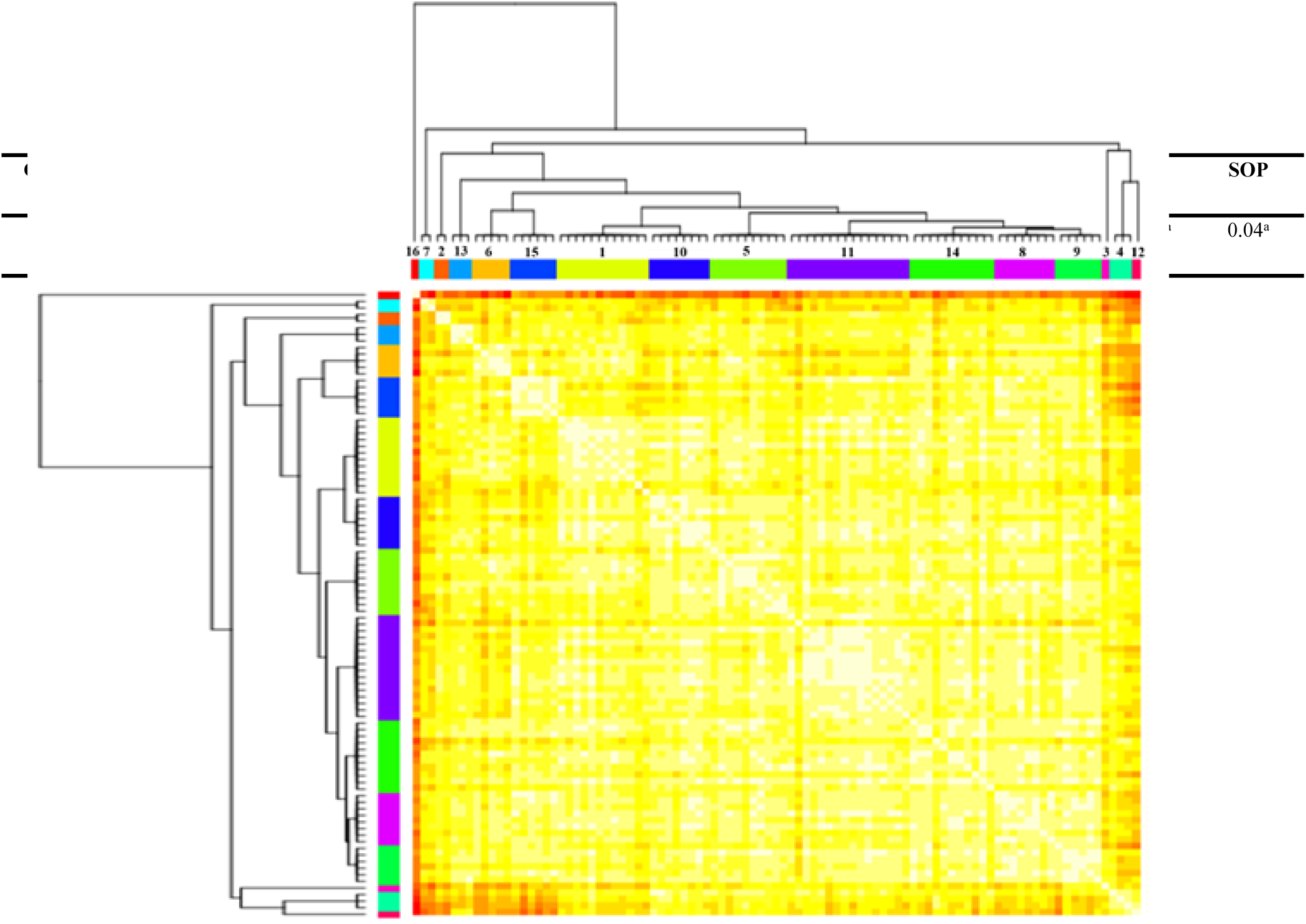
Heatmap and hierarchical clustering of the genetic distances of the *C. moschata* accessions, based on agro-morphological traits, the total carotenoid content of the fruit pulp, and the yields of seeds and seed oil. The coloured bars on the upper and lower axis correspond to the groups obtained in the clustering. Orange colour indicates higher dissimilarity and white indicates lower dissimilarity.

### Identification of promising clusters and *per se* identification of promising genotypes

In order to facilitate the visualization of clusters with the most desirable means for each characteristic, a grouping of means of clusters was performed by the Tocher method (Table 5).

**Table 5.**
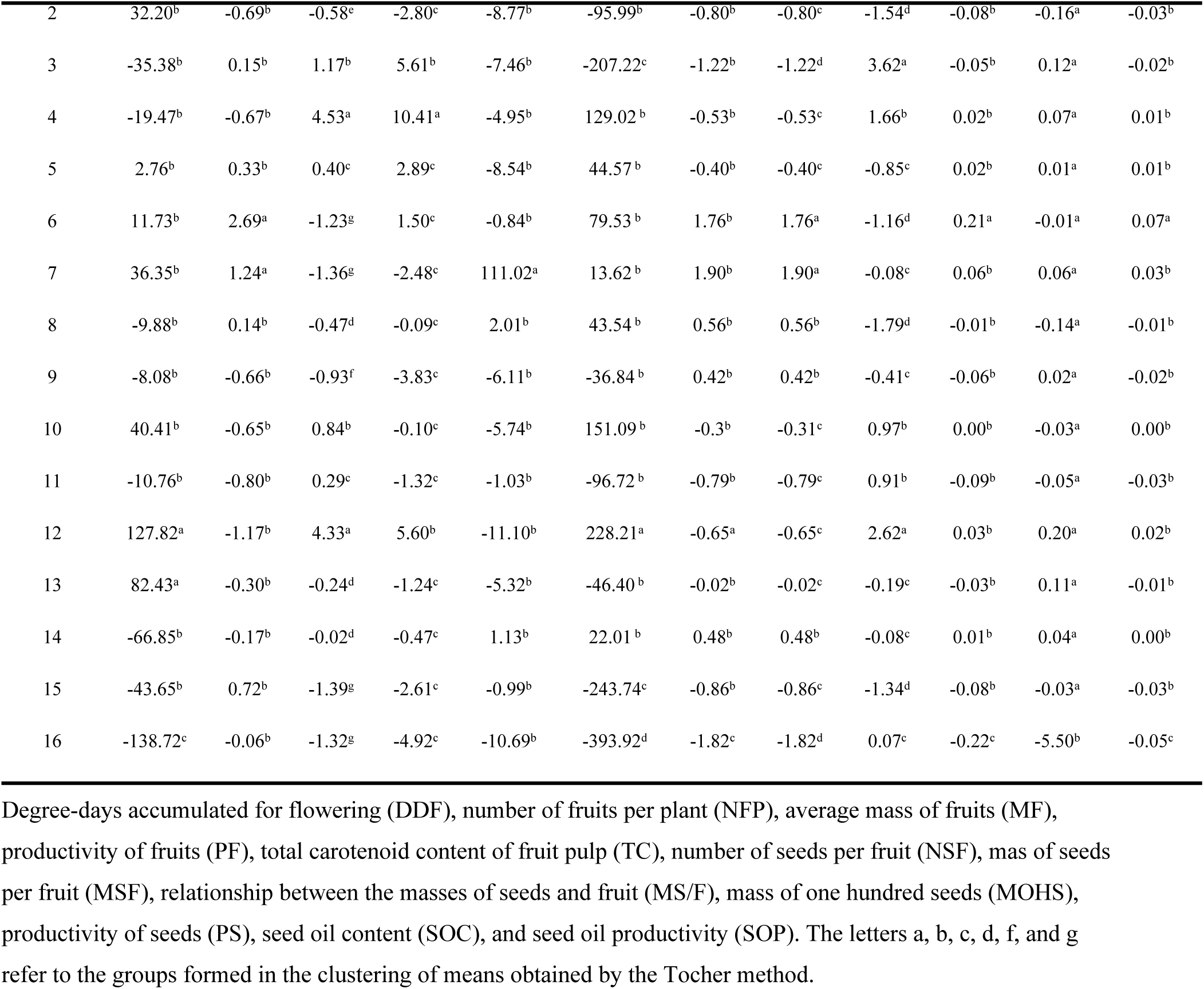
Grouping of means of the genotypic values of the groups obtained in the analysis of variability for agro-morphological aspects, the total carotenoid content of the fruit pulp and the yields of seed and seed oil

The lowest mean for DDF corresponded to group 16, which contained only the control Tetsukabuto, although most groups expressed intermediate averages for this characteristic (Table 5). The group with the highest mean for PF was group 4, formed by the accessions BGH-1927, BGH-4681A and BGH-5653. This group also expressed one of the highest averages for MF and intermediate averages for NFP. As for the TC, the highest average corresponded to group 7, formed by the accessions BGH-5455A and BGH-5598A. Regarding seed and seed oil productivity, it was observed that groups 1 and 6 expressed the highest averages. This first one contained the largest number of accessions (Table 4).

The identification *per se* of the most promising accessions for each trait, based on their respective genotypic effects, is shown in tables 6 and 7. Also in these tables are the estimates, for each accession, of their genetic gains and the new predicted average for each trait.

**Table 6.** Estimates of the genotypic effects, genetic gain and new predicted averages for the degree-days accumulated for flowering (DDF), fruit productivity (PF) and total carotenoid content of the fruit pulp (TC), for the top 15% most promising accessions and the controls

| DDF |  |  |  | PF |  |  |  | TC |  |  |  |
| --- | --- | --- | --- | --- | --- | --- | --- | --- | --- | --- | --- |
| Acessions | g | Gain | New Average | Acessions | g | Gain | New Average | Acessions | g | Gain | New Average |
| BGH-6749 | -291.35 | -355.55 | 251.09 | BGH-4453 | 17.32 | 16.32 | 29.27 | BGH-5455A | 113.85 | 113.70 | 179.46 |
| BGH-5639 | -152.11 | -216.29 | 390.35 | BGH-5653 | 5.60 | 15.44 | 28.38 | BGH-5598A | 108.19 | 108.03 | 173.80 |
| BGH-291 | -119.80 | -183.97 | 422.67 | BGH-5544A | 13.82 | 12.82 | 25.76 | BGH-1461A | 14.18 | 14.03 | 79.80 |
| BGH-5638 | -102.40 | -166.57 | 440.07 | BGH-4681A | 11.74 | 10.74 | 23.69 | BGH-5616A | 11.50 | 11.35 | 77.12 |
| BGH-6587A | -91.41 | -155.57 | 451.07 | BGH-5224A | 10.21 | 9.21 | 22.16 | BGH-6794 | 11.08 | 10.93 | 76.70 |
| BGH-5624A | -83.56 | -147.73 | 458.91 | BGH-6587A | -4.26 | 8.35 | 21.29 | BGH-5556A | 9.85 | 9.70 | 75.47 |
| BGH-1004 | -83.32 | -147.48 | 459.16 | BGH-4590A | 5.61 | 4.61 | 17.55 | BGH-5606A | 8.78 | 8.63 | 74.40 |
| BGH-1749 | -76.12 | -140.28 | 466.36 | BGH-5649 | 5.60 | 4.59 | 17.54 | BGH-5497 | 8.73 | 8.58 | 74.34 |
| BGH-5301 | -76.12 | -140.28 | 466.36 | BGH-5051 | 5.30 | 4.30 | 17.25 | BGH-5451 | 8.68 | 8.53 | 74.29 |
| BGH-5456A | -75.88 | -140.04 | 466.60 | BGH-5596A | 4.52 | 3.52 | 16.46 | BGH-5493A | 8.62 | 8.47 | 74.24 |
| BGH-5485A | -75.88 | -140.04 | 466.60 | BGH-5248 | 4.09 | 3.09 | 16.03 | BGH-5247A | 7.34 | 7.19 | 72.95 |
| BGH-5530A | -75.88 | -140.04 | 466.60 | BGH-5472A | 4.06 | 3.06 | 16.00 | BGH-6749 | 6.91 | 6.76 | 72.53 |
| BGH-6794 | -68.94 | -133.11 | 473.53 | BGH-5473A | 3.62 | 2.62 | 15.57 | BGH-6587A | 13.49 | 6.71 | 72.47 |
| BGH-4598A | -68.09 | -132.25 | 474.39 | BGH-5556A | 3.54 | 2.54 | 15.49 | BGH-95 | 6.73 | 6.58 | 72.34 |
| Average |  |  | 606.64 | Average |  |  | 12.95 | Average |  |  | 65.76 |
| Controls |  |  |  | Controls |  |  |  | Controls |  |  |  |
|  | BLUES | Gain | New Average |  | BLUES | Gain | New Average |  | BLUES | Gain | New Average |
| Jabras | -192.84 | -54.12 | 413.99 | Jabras | 8.53 | 8.87 | 20.88 | Jabras | 14.93 | 16.19 | 80.68 |
| Tetsukabuto | -138.72 | 0.00 | 468.11 | Tetsukabuto | -1.03 | -0.69 | 11.31 | Tetsukabuto | 0.23 | 1.49 | 65.98 |
| Maranhão | 3.99 | 142.71 | 610.82 | Maranhão | -4.60 | -4.26 | 7.75 | Maranhão | -5.13 | -3.88 | 60.61 |
| Jacarezinho | 5.89 | 144.61 | 612.72 | Jacarezinho | -4.92 | -4.57 | 7.43 | Jacarezinho | -10.69 | -9.43 | 55.06 |
| Average |  |  | 526.41 | Average |  |  | 11.85 | Average |  |  | 65.58 |

**Table 7.**
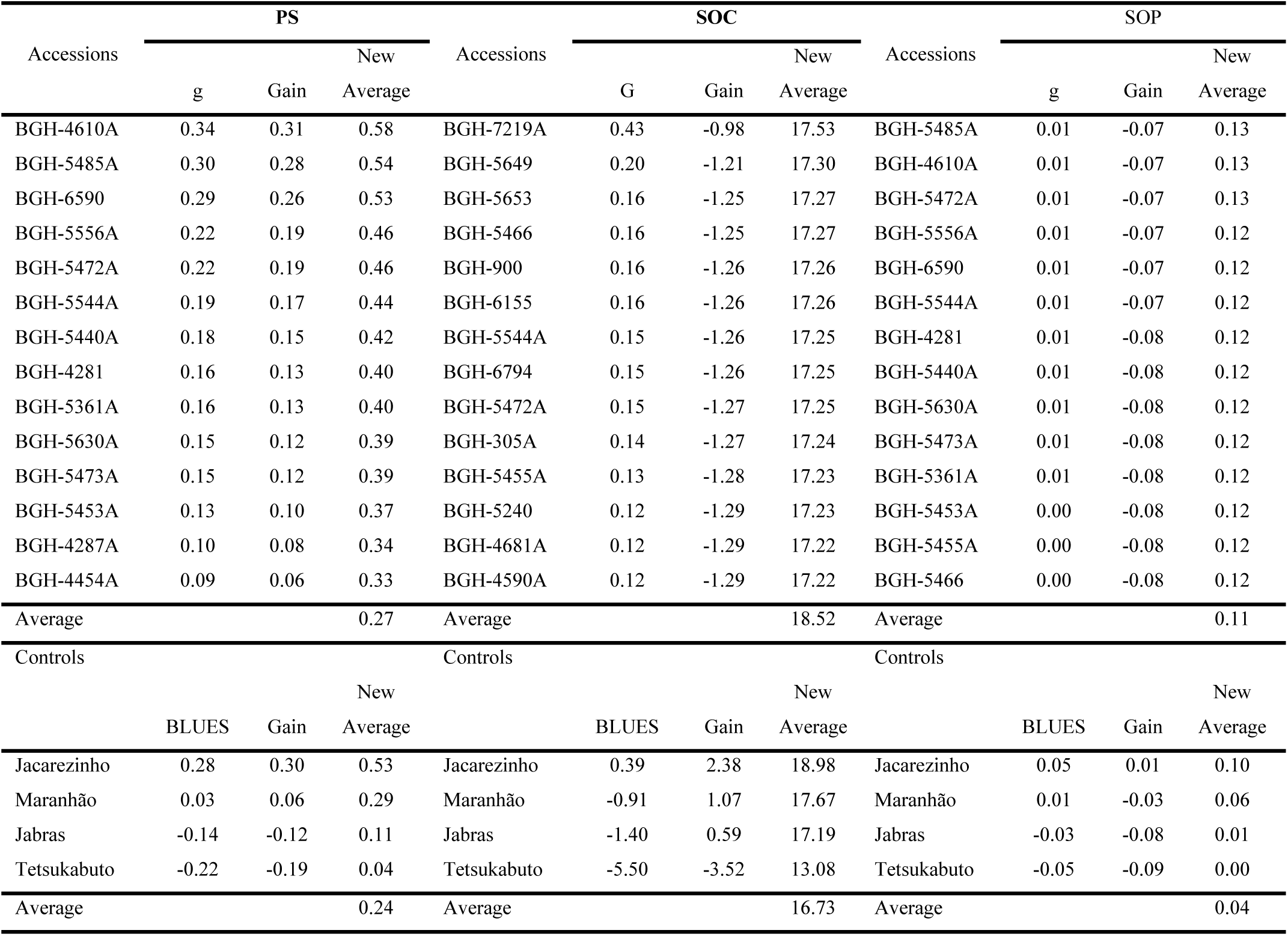
Estimates of the genotypic effects, genetic gain and new predicted averages for the productivity of seeds (PS), seed oil content (SOC), and seed oil productivity (SOP), for the top 15% most promising accessions and the controls

For DDF, the selected accessions expressed averages much lower than the general average of the accessions (606.64) and the average of the controls (526.41). The new predicted averages for this trait among the selected accessions ranged from 474.39 to 251.09, and genetic gains from - 132.25 to -355.55. Notably, the accessions BGH-6749, BGH-5639, and BGH-2191 were the most promising for DDF (Table 6).

For PF, the selected accessions expressed averages higher than the general average of the accessions (12.95 t ha^-1^) and the average of the controls (11.85 t ha^-1^). The new predicted averages for this characteristic among the selected accessions ranged from 15.49 to 29.27 t ha^-1^. As for TC, the selected accessions also expressed averages much higher than the general average of the accessions (65.76 µg g^-1^ of fresh weight) and that of the controls (65.58 µg g^-1^ of fresh weight). The new averages predicted for this characteristic among those selected ranged from 72.34 to 179.46 μg g^-1^ of fresh pulp mass, and the most promising accessions for this characteristic were BGH-5455A and BGH-5598A (Table 6).

The identification *per se* of the most promising accessions for seed productivity (PS), seed oil content (SOC) and seed oil productivity (SOP), together with their respective genetic gains and new predicted averages for these characteristics is shown in table 7.

As for PS, the new predicted averages for this trait among the selected accessions ranged from 0.33 to 0.58 t ha^-1^ and the genetic gains from 0.06 to 0.31 t ha^-1^. Notably, the accessions BGH-4610A, BGH-5485A, and BGH-6590 were the most promising for this characteristic (Table 7). The selected accessions displayed small differences between them for the SOC, however, the average of these was higher than that of the controls (16.73%). Finally, for the SOP trait, the new predicted averages for this trait ranged from 0.12 to 0.13 t ha^-1^ and the genetic gains from -0.07 to -0.08 t ha^-1^. The accessions BGH-5485A, BGH-4610A, and BGH-5472A were the most promising for this characteristic (Table 7).

## DISCUSSION

### Genetic-statistical parameters

As with other species, the usefulness of *C. moschata* germplasm conserved in banks depends on the level and quality of information associated with it [44, 14, 29, 30, 31]. The samples of *C. moschata* maintained by BGH-UFV constitute one of the largest collections of this species in Brazil [32]. Studies involving the assessment of this germplasm have allowed the identification of accessions with crucial characteristics for this crop, such as phytopathogenic resistance, and for its genetic improvement in terms of the productive and nutritional aspects of its fruits and seed oil [34, 20, 12, 35]. Although BGH-UFV maintains more than 350 accessions of *Cucurbita* ssp. [33], part of this germplasm has not yet been assessed, demonstrating the importance of continuing these studies.

Most of the *C. moschata* germplasm shows vigorous growth and indeterminate growth habit [45]. Thus, *C. moschata* plants commonly occupy a large areas of cultivated land, making it difficult to evaluate their germplasm in experimental designs, such as in randomised blocks. The difficulty in ensuring satisfactory homogeneity throughout the experimental area is the main limitation in the evaluation of *C. moschata* germplasm in randomised blocks. In addition, the germplasm seed samples kept in banks are small, making it impossible to repeat accessions throughout the experimental area and assess quantitative characteristics. In view of this, we proposed in this study to evaluate part of the *C. moschata* germplasm maintained at BGH-UFV using the design known as Federer’s augmented blocks [37]. The details of all aspects inherent to this design are very well described by Federer and, according to him, the design circumvents the limitations mentioned above and can be adopted even when the propagating material is insufficient for the establishment of more than one plot and where the quantity of samples to be evaluated is too great.

The present study describes the evaluation of one of the largest germplasm volumes of *C. moschata*. The germplasm expressed markedly high genotypic variances for characteristics related to seed production such as NSF and MSF, for DDF, and for TC and PF (Table 3). Most of the phenotypic variance estimates of these characteristics were due to the contribution of genotypic variance. These results corroborate those reported by [46], who also observed higher estimates of genotypic variances for NSF and flowering characteristics, and also a greater contribution of genotypic variance to the phenotypic variance in these characteristics. Additionally, most of the characteristics expressed high estimates of heritability (>0.50), according to the classification of [47]. These estimates were very high for the characteristics of seeds such as MSF, MS/F, and NSF, and for aspects related to fruits, such as TC and MF. High estimates of heritability point to a greater correlation between the phenotype and the genotype [48], indicating that most of the variability observed for these characteristics resulted from genotypic effects.

The high estimates of genotypic variances may be associated with the quantitative nature of these characteristics, which may be the result of the influence of a high number of genes [49]. Most of the germplasm evaluated in this study came from the land of family-based farmers, who do not carry out selection for seed characteristics, nor with a view to obtaining earlier-flowering genotypes. As already mentioned, the exchange of seeds between farmers and the natural occurrence of hybridization between populations of *C. moschata*, has increased the variability of this species, even for characteristics for which selection is commonly carried out, such as fruit productivity.

The high estimates of genotypic variance and heritability allowed considerable predicted gains through selection for most of the characteristics (Table 3). For the number of degree-days accumulated for flowering, a gain of -92.947 was obtained, a considerable result, taking into account the average of the evaluated accessions (606.642). It was possible to obtain a gain of 7.817 t ha^-1^ for PF and 20. 426 μg g^-1^ of fresh pulp mass for TC. The gains for PS and SOP were 0.187 and 0.072 t ha^-1^, respectively (Table 3).

The average relationship between the coefficient of genetic variation and the residual coefficient was close to one unit for most of the characteristics. Although the estimates of the residual coefficient of variation for most characteristics were high, in general they tended to be lower in relation to their corresponding coefficient of genotypic variability, which demonstrates that most of the variability expressed by germplasm was due to genetic factors (Table 3).

### Genetic correlation network

Analysis of correlations between characteristics has been widely used in plant breeding, where often a high number of characteristics must be considered simultaneously [50, 51]. As for PF, a positive correlation was observed with MF. Fruit productivity also expressed positive and high correlations with aspects related to fruit dimensions such as DIC, HF and DF (Figure 1). The correlation between NFP and PF was 0.39 and although the estimates of correlations between PF and the other characteristics of this group were low (<0.70), all of them were significant (*p*<0.001).

Correlation analysis is often used to assist in indirect selection for certain characteristics [52, 51]. However, as highlighted by [53], in cases where there is an intention to practice indirect selection for a primary characteristic by means of a secondary one, it is necessary that the heritability of the latter characteristic be greater than that of the former so that the selection is efficient. In view of this, the selection of genotypes with higher MF seems to be a promising alternative for obtaining higher fruit productivity in *C. moschata*. It should, however, be highlighted that when selecting genotypes with the aim of increasing fruit productivity in *C. moschata*, crucial aspects for their acceptability in the consumer market, such as the shape and size of fruits, must be considered. Currently, important pumpkin consumption centres like the state of Minas Gerais and most of the southeast region of Brazil demand smaller fruits, and most of the consumption in these regions is represented by fruits from hybrid cultivars, such as Jabras and Tetsukabuto, which have a globular format and weight ranging from 2 to 3 kg [14]. On the other hand, in the north and northeast regions of Brazil, there is greater acceptability for larger fruits, which are commonly sold in slices. The prevention of waste and the ease of transport are the determining aspects for the acceptability of fruit patterns, and the search for greater productivity in the cultivation of *C. moschata* must therefore also consider these characteristics, equating them with aspects such as the NFP, HF and DF.

Another correlation group, the discussion of which is relevant in terms of cohesion, was that formed by PS and characteristics such as MS/F, NSF and MSF. As can be seen in figure 1, the correlations of PS with these characteristics were positive. It should also be noted that the NFP and PF expressed high correlations with PS (0.74 and 0.51, respectively), and all of them were significant (*p*<0.001).

Based on these results, the simultaneous consideration of aspects such as higher NFP, higher PF and higher MS/F relationship seems to be a promising alternative for obtaining higher seed productivity in *C. moschata*. The heritability estimates obtained for these characteristics (>0.42), suggest the feasibility of reasonable gains with the selection for each one of them (Table 3). As mentioned previously, PF and NFP displayed a correlation of 0.39. The general average of the accessions for this first trait was 12.94 t ha^-1^, and some of them expressed new predicted averages of up to 29.27 t ha^-1^, in the case of BGH-4453 (Table 6). The accessions BGH-5653, BGH-5544A, BGH-4681A, BGH-5224A, and BGH-6587A all displayed productivities greater than 20 t ha^-1^. With this, besides greater PF and NFP, the selection of genotypes with higher PS should also prioritise greater translocation of photoassimilates for seed production, something indicated by a higher MS/F ratio.

Despite its applicability, the analysis of correlation has some limitations, and, as warned by [54] the quantification and interpretation of the correlation coefficients between two or more characteristics can result in errors during the selection process. According to them, this occurs because the high estimates of correlations between these characteristics may be the result of the effect of one or more secondary characteristics. It is therefore recommended that the analysis of the association between a primary and secondary characteristic be accompanied by information on the direct and indirect effects of the secondary variables on the primary [55], an approach currently known as path analysis [54].

Despite some limitations, correlation analysis has proven to be quite useful in plant breeding, mainly in the indirect selection for one or more main characteristics with low heritability or of difficult assessment. This indirect selection is carried out based on secondary characteristics, with greater heritability or the assessment of which is easier, providing faster genetic gains compared to direct selection. In fact, correlation analysis has assisted in the indirect selection for characteristics of roots [56], for productivity in different crops [57, 58, 59], and for nutritional aspects and quality of fruits [60, 61]. Correlation analysis can also be very useful in the characterization and management of plant germplasm, as it has the ability to optimise the choice and the number of descriptors to be used in this process.

### Genetic variability and clustering

The analysis of variability provides important assistance in the initial phase of plant breeding programs and in the management of plant germplasm. In this first case, it provides the allocation of accessions in groups, guiding the conduction of crossings. *C. moschata* is allogamous, therefore analysing the variability of its germplasm can assist in the orientation of crossings between more diverse genotypes, thereby aiding the exploration of hybrid vigour [62, 5]. Regarding the assistance in the management of plant germplasm, variability analysis allows the identification of duplicates in the germplasm collections [63, 64, 65], which correspond to pairs or groups of accessions with high similarity. In fact, it is estimated that less than 30% of the accessions maintained in the collections worldwide are distinct, which hinders their maintenance [28]. Therefore, in addition to optimizing the use of germplasm, the variability analysis reduces the cost of its maintenance by reducing its volume [66].

The accessions of *C. moschata* assessed in this study displayed high genetic variability in their agro-morphological characteristics, the total carotenoid content of the fruit pulp, and the productivity of seeds and seed oil, resulting in the formation of 16 clusters (Table 4). There was low similarity between the clusters formed, as shown by the predominance of yellow colour in the hierarchical clustering in heatmap format (Figure 2). The visual analysis of this cluster also indicates the homogeneity of the genetic distances between clusters, which verified the clustering efficiency. As can also be seen in figure 2, there was uniformity in the yellow colour for the genetic distances between groups, confirming the homogeneity of distances between them. The variability denoted by the clustering of the accessions corroborates the high estimates of genetic variances and heritabilities displayed by most of the agronomic characteristics, the total carotenoid content of the fruit pulp, and the seed characteristics such as MSF, MS/F and NSF (Table 3). This is also analogous to other studies involving the analysis of variability in this crop in Brazil [18, 20].

### Identification of promising groups of genotypes

The analysis of averages of the groups using the Tocher method (Table 5) provided information on the similarity or divergence between the groups, allowing the identification of those with more desirable averages for each characteristic. In *C. moschata*, this approach can assist in the orientation of crossings targeting hybrid vigour exploitation and the segregation of populations for their characteristics of interest [67, 68].

Group 11 contained the largest number of clustered accessions, 15 in total, together with Jabras, one of the controls. Group 1, the second largest, contained 10 accessions and two controls (Jacarezinho and Maranhão). The clustering of these two cultivars with similar characteristics in the same group reflects the clustering consistency. As shown in table 5, this group expressed a high genotypic average for TC and the highest averages for PS and PCOS, confirming the high number of promising accessions for these characteristics. Groups 5 and 14 contained 10 and 11 accessions, respectively, making them the next largest groups formed.

Regarding TC, the highest average was in group 7, formed by the accessions BGH-5455A and BGH-5598A (Table 4). These accessions were also identified as the most promising for TC in the identification *per se*, with new predicted averages greater than 170 μg g^-1^ of fresh pulp mass (Table 6). This result is much higher than those reported by previous studies [69, 35, 6]. Among these, the study of [35], for example, involving the characterization of 55 accessions of *C. moschata*, also maintained by the BGH-UFV, reported TC averages not greater than 118,70 μg g^-1^ of fresh pulp mass. On the other hand, [1] and [70] reported TC averages of up to 404.98 μg g^-1^ of fresh mass, when evaluating *C. moschata* germplasm from northeast Brazil. The differences observed for TC between the present study and previous studies might be mainly associated with the genetic makeup of the germplasm evaluated in each study. According to [70], in the northeast region of Brazil there is a preference for winter squash fruits with more orange pulp, a characteristic associated with higher levels of carotenoids, which corroborates the results obtained for this characteristic in studies involving the evaluation of *C. moschata* germplasm from this region.

Studies with *C. moschata* commonly involve the analysis of fruit pulp carotenoids and generally report high levels of these components [1, 71, 72, 6]. Among these studies, [1] reported the identification of about 19 different carotenoids in the carotenogenic profile of the fruit pulp, and found that *β*- and *α*-carotene constitute the largest proportion of the total carotenoid content in this species. In fact, this vegetable has been considered one of the best sources of carotenoids such as *β*-carotene, with levels above those found in other important carotenogenic vegetables, such as carrots [73].

The main biological functions of components such as *α*- and *β*-carotene are their pronounced pro-vitamin A activity [74, 75], and a series of bioactive functions, especially antioxidant activity [76, 77]. Along with its bioactive functions, *C. moschata* brings together fundamental characteristics for biofortification programmes, such as high production potentials and profitability, high efficiency in reducing deficiencies in micronutrients in humans, and good acceptance by producers and consumers in the growing regions [10]. *C. moschata* has therefore been strategically used in programmes targeting biofortification in vitamin A precursors, among them the Brazilian Biofortification Programme (BioFORT), led by Embrapa [11].

Regarding PS and SOP, the main interest in the assessment of these traits in *C. moschata* corresponds to the high potential of using oil from its seeds for food purposes. This vegetable has a high oil content, with the lipid fraction of its seeds reaching up to 49% of its composition [78]. The lipid profile of this oil consists of more than 70% unsaturated fatty acids, with a preponderance of fatty acids such as linoleic C18: 2 (Δ^9,12^) and oleic C18: 1 (Δ^9^). In this regard, there is an interest among governments and health experts in encouraging the consumption of unsaturated fatty acids rather than saturated ones, based on the consensus that this reduces the risk of cardiovascular diseases [79, 80, 81].

*C. moschata* seed oil is also rich in bioactive components such as vitamin E and carotenoids [4, which have important antioxidant activity, in addition to providing protection to the oil during its conservation. Despite this, most of the seeds from the production of *C. moschata* in Brazil are still discarded during consumption. Their use therefore represents an alternative for supplementing the diet and increasing the income of farmers involved in the production of this vegetable.

Group 16, formed solely by the control Tetsukabuto, displayed the lowest average DDF (Table 5), indicating that this genotype has the earliest flowering period. As can also be seen in this table, most groups expressed intermediate averages for DDF. Normally, *C. moschata* plants have very long internodes, and this, coupled with the vigorous growth of this species, represents a limitation on its cultivation since plants with a greater internode length require much larger areas for cultivation. The interest in assessing precocity in *C. moschata* is based on the possible relationship of this characteristic with the development of shorter vines or the habit of determined growth. According to [82], the *Bu* gene, identified as being responsible for the formation of shorter internodes in pumpkins, is also linked to earlier flowering in this species. In a study involving the evaluation of hybrids and segregating winter squash populations for oil production and plant size reduction, [43] observed that the cultivars Piramoita and Tronco Verde, which have determined growth habits, displayed the smallest number of days for female flowering. Greater precocity is an important characteristic for most crops, especially in the cultivation of vegetables. This feature optimises the use of cultivation areas, reduces the risks of exposure of the crop to adverse abiotic and biotic factors, and reduces management costs.

Group 4, formed by BGH-1927, BGH-4681A and BGH-5653, expressed the highest average PF (Table 5). This group also expressed one of the highest averages for MF and an intermediate average for NFP, corroborating the estimates of the correlations between these characteristics and PF (Figure 1). The accessions BGH-4681A and BGH-5653 were also identified as the most promising for PF in the *per se* identification, with averages above 20 t ha^-1^ (Table 6). These averages were much higher than the world average, estimated at 13.4 t ha^-1^ [8].

Although the cultivation of *C. moschata* is primarily intended for fruit production, as already mentioned, the selection of genotypes for greater fruit productivity in this crop must also consider crucial aspects for the acceptability of fruits such as shape and size. In general, winter squash production must currently prioritise the adoption of cultivars with smaller fruits. In addition to obtaining fruits of greater mass, greater productivity in *C. moschata* can also be achieved by obtaining cultivars with higher NFP, based on the estimated correlation observed between PF and NFP (Figure 1).

### *Per se* identification of promising accessions

*Per se* identification of the most promising accessions for the characteristics considered crucial in the production of *C. moschata* is shown in tables 6 and 7. This approach can guide selection for a specific trait, allowing the identification of promising accessions for the development of superior inbred lines and/or open-pollinated cultivars. In fact, from a brief survey of the Brazilian National Cultivar Register (RNC), it appears that of the 182 cultivars of *C. moschata* registered at the moment, most of them consist of open-pollinated cultivars [83]. This survey, also found a considerable number of intra- and interspecific hybrids, confirming the feasibility of applying inbreeding in certain stages of *C. moschata* breeding.

For DDF, the selected accessions displayed averages much lower than the general average of the accessions (606.64 degree-days) and the controls (526.21 degree-days). Notably, the accessions BGH-6749, BGH-5639, and BGH-219 expressed the lowest new predicted averages for DDF, making them the earliest-flowering accessions (Table 6). Regarding PF, the notably more promising accessions were BGH-4453, BGH-5653, BGH-5544A, BGH-4681A, BGH-5224A, and BGH-6587A, which expressed gains and new predicted averages for fruit productivity above 8 and 20 t ha^-1^, respectively (Table 6). As can also be seen in this table, these accessions displayed gains and new predicted averages much higher than those of the controls. It should be highlighted that the BGH-5544A accession also expressed high averages for PS and SOP, corroborating the correlations of these characteristics with PF (Figure 1). This indicates the potential for the dual use of this accession to produce fruit and seed oil.

Regarding TC, the most promising accessions were BGH-5455A and BGH-5598A (Table 6). These accessions expressed gains and new predicted averages for TC higher than 108.03 and 173.80 μg g^-1^ of fresh pulp mass, respectively, much higher than those of the controls. For the characteristics of seed and seed oil, it was found that the accessions BGH-4610A, BGH-5485A, and BGH-6590 were the most promising for PS (Table 7). These accessions expressed gains and new predicted averages for seed productivity of up to 0.31 and 0.58 t ha^-1^, respectively. The most promising accessions for SOP were BGH-5485A, BGH-4610A, and BGH-5472A, which expressed new predicted averages for seed productivity of 0.13 t ha^-1^. It is worth highlighting that these accessions corresponded to those with higher PS, corroborating the strong correlation between PS and SOP (Figure 1).

## CONCLUSIONS

The accessions of *C. moschata* assessed in this study expressed high genetic variability for agro-morphological characteristics and for agronomic aspects related to the production of seeds such as NSF and MSF, for DDF, and for TC and PF, which allowed the obtainment of considerable gains from selection for each of these characteristics.

The network of genetic correlations showed that higher fruit productivity in *C. moschata* might be achieved from the selection of aspects considered crucial in the production of this crop such as higher NFP, HF and DF. It also showed that greater seed productivity might be achieved with the selection for higher MS/F, NSF and MSF; information that will assist in selection for higher productivity of fruit, seed and seed oil.

The clustering analysis resulted in the formation of 16 groups, with low similarity between the groups, which corroborates the variability of these accessions.

The grouping of the averages of the clusters and the identification *per se* allowed the recognition of the most promising groups and accessions for each characteristic, an approach that will guide the use of these accessions in breeding programs.

*Per se* analysis identified the accessions BGH-6749, BGH-5639, and BGH-219 as those with the lowest averages for DDF, highlighting them as the earliest flowering accessions. The most promising accessions for PF were BGH-4453, BGH-5653, BGH-5544A, BGH-4681A, BGH-5224A, and BGH-6587A, with new predicted averages greater than 20 t ha^-1^. The accessions with the highest averages for TC were BGH-5455A and BGH-5598A, with averages greater than 170.00 μg g^-1^ of fresh pulp mass. The accessions BGH-5485A, BGH-4610A, and BGH-5472A were the most promising for SOP, also corresponding, in the case of the former two, to those with the highest averages for PS. The accessions of *C. moschata* assessed in this study are a promising source for the genetic improvement of characteristics such as early flowering, total carotenoid content of the fruit pulp, and productivity of seeds and seed oil.

## ACKNOWLEDGMENTS

We are thankful to the Coordenação de Aperfeiçoamento de Pessoal e Nível Superior - Brasil (CAPES) - Finance Code 001. We are also thankful to the National Council for Scientific and Technological Development–CNPq, for the additional financial support in this study.

## AUTHOR CONTRIBUTIONS

The authors declare that they contributed according to the specification below.

**Ronaldo Silva Gomes**: Investigation, data curation, writing and editing of the original draft.

**Ronaldo Machado Junior, Cleverson Freitas de Almeida, Rebeca Lourenço de Oliveira, Fabio Teixeira Delazari**: Investigation and data curation.

**Rafael Ravaneli Chagas**: Software.

**Derly José Henriques da Silva**: Conceptualization, supervision, and Writing – review & editing of the original draft.

